# Individuals learning to drive solo before age 18 have superior spatial navigation ability compared with those who learn later

**DOI:** 10.1101/2023.09.26.559011

**Authors:** Emre Yavuz, Ed Manley, Christoffer J. Gahnstrom, Sarah Goodroe, Antoine Coutrot, Michael Hornberger, Hugo Spiers

**Affiliations:** Institute of Behavioural Neuroscience, Department of Experimental Psychology, Division of Psychology and Language Sciences, University College London, London, UK; Faculty of Environment, School of Geography, University of Leeds, Leeds, UK; Department of Psychology, University of Pennsylvania, Pennsylvania, USA; LIRIS - CNRS - University of Lyon, France; Norwich Medical School, University of East Anglia, Norwich, UK

## Abstract

A challenge associated with driving vehicles can be navigating to the destination. While driving experience would seem beneficial for improving navigation skill, it remains unclear how driving experience relates to wayfinding ability. Using the mobile video game-based wayfinding task Sea Hero Quest, which is predictive of real-world navigation, we measured wayfinding ability in US-based participants (n = 694, 291 men, 403 women, mean age = 26.8 years, range = 18-52 years). We also asked travel-related self-report questions, including the age one started learning to drive, the age one started driving solo and weekly driving hours. A multivariate linear regression model found that those who started driving solo below aged 18 had significantly better wayfinding ability than those starting to drive solo aged 18 and above. Other driving-related self-report measures were not associated with wayfinding ability. Future studies should determine the directionality of the association between driving experience and wayfinding ability.

## Introduction

Being able to maintain a sense of direction and location in order to find our way in different environments is a fundamental cognitive function that relies on multiple cognitive domains (Newcombe et al., 2022; Spiers et al., 2023). Human navigation involves a range of processes such as planning routes, reading maps, identifying landmarks and maintaining a sense of direction (Newcombe et al., 2022). Understanding individual differences in navigation ability is crucial, given that deficits in navigation may constitute the earliest signs of Alzheimer’s Disease (Coughlan et al., 2018), and are apparent in other conditions such as Traumatic Brain Injury (Seton et al., 2023). There are also negative effects of disorientation, such as getting lost in everyday environments, leading to distress for patients and family members and in extreme cases death from exposure (Coughlan et al., 2018). Understanding individual differences in navigation will also advance the field of spatial cognition at large (Newcombe et al., 2022; Spiers et al., 2023). Creating a valid test of navigation that accounts for the wide variation in navigation ability is challenging, given the large sample sizes needed and high levels of environmental manipulation and experimental control required in standard research settings (Newcombe et al., 2022). With the recent evolution of widespread touch-screen technology on both tablet and mobile devices and virtual reality (VR), our team developed a series of navigation tests in the form of a mobile video game app Sea Hero Quest (SHQ)(Spiers et al., 2023). SHQ has since been used to test the navigation ability of 4 million people globally, has good test-retest reliability and is predictive of real-world navigation ability (Coughlan et al., 2020; Coutrot et al., 2018, 2019).

Many studies have shown that spatial cognition is associated with driving ability, where poorer performance on various facets of spatial cognition, including spatial perspective taking, mental rotation and visuospatial processing performance, is associated with worse driving ability (Andrews & Westerman, 2012; Di Meco et al., 2021; Kunishige et al., 2020; Moran et al., 2020; Tinella et al., 2020, 2021a, 2021b). Despite these associations between driving ability and spatial cognition, it remains unclear how driving experience per se is associated with spatial cognition. Additionally, no studies have investigated how driving experience relates to wayfinding in particular, a key facet of spatial cognition required for successful spatial navigation where one must actively navigate to a goal from a start location using a representation of their environment in the form of a mental map (Patai & Spiers, 2021). Both poorer driving and wayfinding ability are associated with greater risk of cognitive decline and neurodegenerative disease (Coughlan et al., 2019; Duchek et al., 2003; Ott et al., 2008; Roe et al., 2016; Uc et al., 2017), where current drivers have significantly slower cognitive decline than former drivers (Choi et al., 2013). Examining the association between driving experience and wayfinding ability may therefore provide a better understanding as to why certain individuals are at greater risk of cognitive decline.

Given the many aforementioned studies showing significant positive associations between driving ability and spatial cognition, we hypothesised that those with greater driving experience, defined by greater hours of weekly driving, a younger age at which one learns to drive and a younger age at one starts to drive solo, would have better wayfinding ability. We also included self-reported measures of GPS use, video games experience, and other related variables as covariates in our analysis because of their potential relation to wayfinding ability (see Yavuz et al., 2023a).

## Methods

### Participants

903 participants living in the US aged 18 and above (395 men, 489 women, mean age = 27 years, SD = 8.0 years, range = 18-66 years, mean number of years of formal education = 16.1 years, 315 currently living in cities, 588 currently living outside cities) were recruited using the Prolific database (www.prolific.co, 2023) and reimbursed for their time. Ethical approval was obtained from the University College London Review Board conforming to Helsinki’s Declaration. All participants provided informed written consent. We first removed 94 participants who reported that they did not currently drive (12% of the sample), and then removed an additional 17 participants who selected ‘other’ for gender given that there were too few participants in this category to run the mediation analysis. We next identified outliers using Mahalanobis’ Distance, a method shown to have high sensitivity and specificity and a minimal change in bias in simulated and real datasets based on questionnaire data when removing outliers compared to other methods (Curran, 2016; Zijlstra et al., 2011)(Supplementary Materials). 97 participants were removed as outliers. This resulted in a final sample size of 694 participants (291 men, 403 women, mean age = 26.8 years, SD = 7.4 years, range = 18-52 years, mean number of years of formal education = 16.2 years, 232 currently living in cities, 462 currently living outside cities). Demographic information for the final sample is summarised in Table 1. Data analysis was completed using R studio (version 1.4.1564) and Python (version 3.9.12).

**Table 1.** Summary of basic descriptive statistics for the variables included in the model (weighted wayfinding distance, age, weekly hours of video game use on all devices, weekly hours of phone use) for men and women separately, and across gender. Mean and standard deviation (SD) values are shown.

|  | **Women** | **Men** |
| --- | --- | --- |
| n (%) | 403 | 291 |
| Weighted wayfinding distance | 12.15 (3.66) | 10.70 (3.75) |
| Age | 25.52 (6.93) | 28.53 (7.72) |
| Hours of phone use per week | 35.03 (22.34) | 25.42 (18.70) |
| Weekly hours of video gaming on all devices | 4.33 (4.91) | 9.30 (6.32) |

### Statistical Power Analysis

A power analysis was conducted using G* power (Erdfelder et al., 1996). Given that a previous study looking at the association between spatial mental transformation skills and driving performance found medium-to-large effect sizes (Tinella et al., 2020), it was determined that 694 participants were sufficient to achieve a medium effect size (Cohen’s f2 = 0.15) at an alpha threshold of 0.05 with 95% power (Selya et al., 2012).

### Experimental Procedure

#### Self-report questionnaires

To characterise driving-related activities, participants answered a series of driving-related questions including the number of hours spent driving, the age they learnt to drive and the age they started driving solo. We also asked participants a broader series of questions about travel-related activities, including the hours they spent cycling, biking, boating and taking public transport per week (five separate questions), the hours of weekly exercise they engaged in, the hours of weekly exercise they engaged in within the last 6 months and the mode in which they thought about space (thinking in distance, blocks or time)(Supplementary Materials). The number of hours per week spent biking, boating and taking public transport were converted to binary yes/no variables (weekly biking yes/no, weekly boating yes/no, weekly public transport yes/no), given the very small number of participants who engaged in these activities on a weekly basis.

To characterise reliance on GPS, the GPS reliance scale was used (Dahmani and Bobot, 2020). This scale has seven items and assesses the frequency at which people have relied on GPS in different situations within the past month (e.g., “How often do you use GPS to travel new routes to a previously visited destination?”). The average score across questions was calculated for each participant. To characterize video game use, participants were asked to indicate the number of hours per week spent playing video games on all devices and the number of hours of phone use per week. Participants were also asked to report their age, gender and highest education level. Please see the Supplementary Materials for the full set of questionnaires.

### Sea Hero Quest Task

Sea Hero Quest (SHQ) is a VR-based video game for mobile and tablet devices which requires participants to navigate through a three-dimensional world on a boat to search for sea creatures in order to photograph them, with the environment consisting of rivers and lakes (Coutrot et al., 2018). Although navigational abilities on SHQ have been assessed using Wayfinding, Path Integration and Radial Arm Maze measures, we focused on wayfinding in this study (Figure 1). Please refer to Yavuz et al., 2023a and 2023b for specific details.

**Figure 1.**
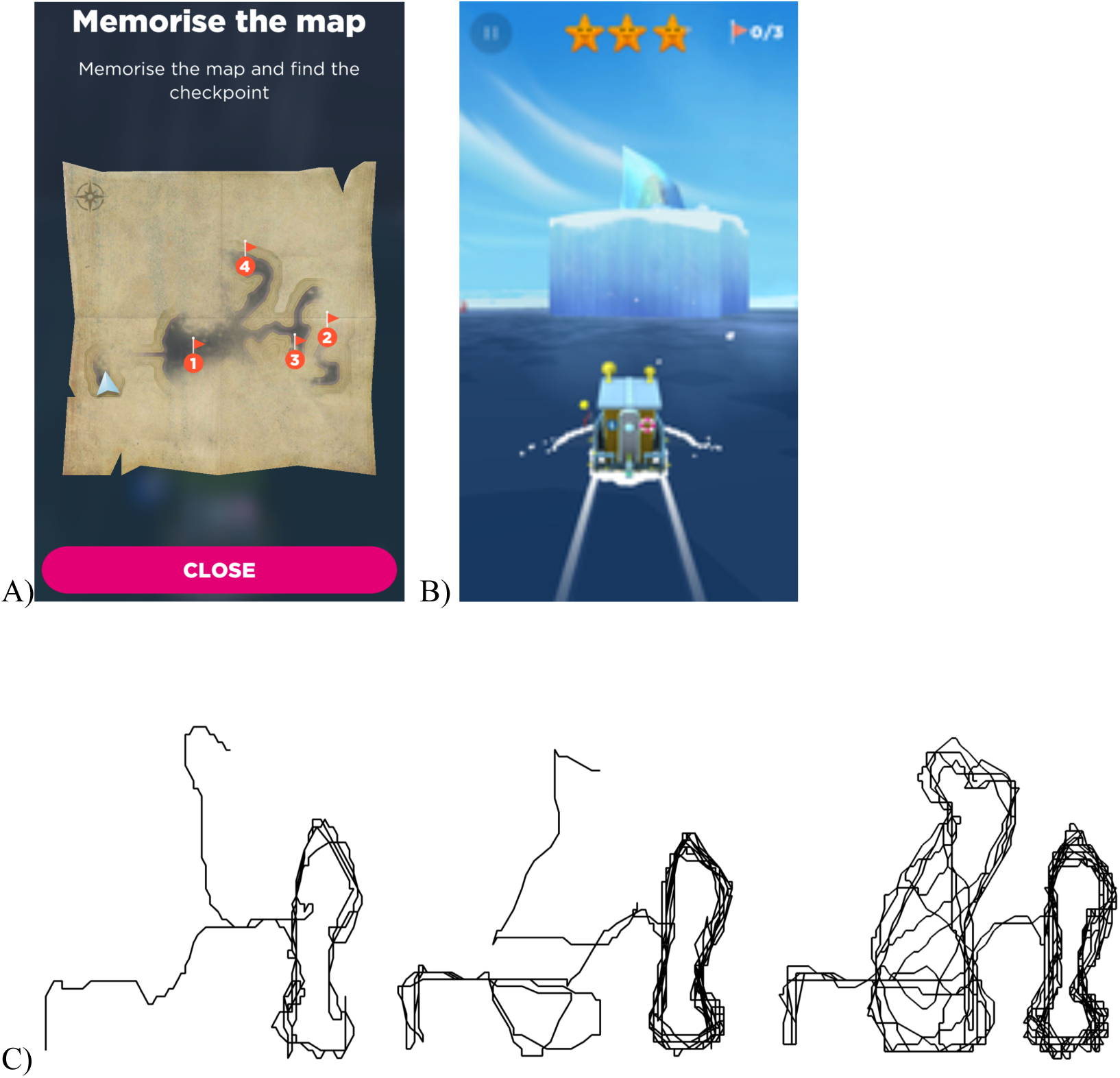
Outline of the wayfinding task. (A) At the start of each level presented participants were presented with a map indicating the goals which they had to navigate to in the order indicated, where ‘1’ indicates goal number 1 etc. (the map from level 68 is shown as an example). Level 1 (not shown) provided one goal and a simple river to reach the goal as training. (B) Participants selected to close the map by pressing ‘close’, at which point the participant had to start navigating to the goals. First-person view of the environment is shown where the participant tapped left or right of the boat to steer it. (C) Examples of individual player trajectories (level 68) from the start location to the final goal. Trajectories are ordered by performance, with the top left providing the best performance (shortest trajectory length), through to bottom right who has the worst performance (longest trajectory length). Adapted from Coutrot et al. (2018).

### Data Analysis

#### Correlations

To measure the independent associations between self-report questionnaire responses and wayfinding distance, we ran a series of correlations. Firstly, Spearman’s correlations were conducted to measure the associations between weighted wayfinding distance and the continuous variables of hours of driving per week, the age one started driving solo, the age one started to learn to drive, hours of walking per week, average GPS reliance score, hours of exercise per week and the hours of exercise per week in the past 6 months. Secondly, point biserial correlations were run to measure the association between the binary categorical variables of engaging in weekly boating (yes/no), engaging in weekly cycling (yes/no), engaging in weekly public transport (yes/no) and gender (man/woman) and weighted wayfinding distance. Thirdly, to measure the associations between weighted wayfinding distance and both 1. The mode of thinking about space (blocks/distance/time) and 2. Hours of daily travel (0-0.5 hours/0.5-1 hours/1-1.5 hours/1.5+ hours), we ran two separate one-way ANOVAs. For Spearman’s correlations, point-biserial correlations and one-way ANOVAs, spearman’s correlation coefficients, point-biserial correlation coefficients and partial-eta squared (*ηp2)* effect sizes were reported respectively (Levine and Hullet, 2002).

#### Multivariate linear regression

We then ran a multivariate linear regression model to analyse the effects of the hours of driving per week, the age one started driving solo and the age one started to learn to drive on wayfinding distance. Covariates included age, gender, highest education level, weekly hours of video gaming on all devices, weekly hours of phone use, hours of daily travel, hours of walking per week, engaging in weekly public transport (yes/no), engaging in weekly boating (yes/no) and engaging in weekly cycling (yes/no), average GPS reliance score and one’s mode of thinking about space (whether they usually thought about space in terms of blocks, distance or time)(Table 4). These covariates were chosen to control for variables that may influence the association between driving-related variables and spatial navigation ability (Ben-Elia, 2021; Coutrot et al., 2022a; 2022b; Jabbari et al., 2022; Sahlqvist et al., 2012; Yavuz et al., 2023). Given that the majority of participants had a daily travel time of 0-0.5 hours per day, we binned daily travel time into two groups before entering this variable into the model: 1. Those travelling for less than 0.5 hours per day and 2. Those travelling for 0.5 hours or greater per day. Additionally, given that our sample was solely based in the US, where most urban environments have a grid-like structure (Coutrot et al., 2022b), we grouped the mode of thinking about space into two groups before entering this variable into the model: 1. Those thinking about space in terms of ‘blocks’ and 2. Those thinking about space in terms of ‘distance’ or ‘time’. Finally, hours of weekly exercise over the last 6 months was binned into two groups before entering this variable into the model: 1. Up to 2 hours per week and 2. 2 hours or more per week.

As a check for multicollinearity, we calculated the variance inflation factor (*VIF*) for each predictor variable in the model (Supplementary Tables S1 and S2). A *VIF* value of <5 indicates that multicollinearity is not a concern (Fox and Monet, 1992; Kim, 2019). Post-hoc t-tests bonferroni-corrected for multiple comparisons to control for type 1 errors were carried out where main effects and interactions were significant. When running post-hoc tests for the driving-related variables (age one started to drive solo and age one started to learn to drive), we binned the data into two categories: <18 and >= 18 years of age, given that 18 years old is the youngest age at which one can obtain a driving license without any imposed driving restrictions (Wang et al., 2020).

#### Mediation analysis

After running the multivariate linear regression model, we ran a mediation analysis to see whether the effect of each of the independent predictor variables that showed a significant association with wayfinding distance was mediated by a. age, b. gender, c. hours per week gaming on all devices, d. whether one grew up in a city or not, given previous associations between age, gender, gaming experience and the environment one grew up in and their wayfinding performance on SHQ (see for more: Coutrot et al., 2018, 2022a, 2022b; Yavuz et al., 2023a, 2023b). We also included the current environment one lives in at the time of the study (city or not) as a mediator for the effect of weekly biking (yes/no) on wayfinding distance, as we were interested in whether the environment one currently lives in would influence the effect of travel-related activities on navigation. Non-parametric bootstrapping with 5000 simulations was used, with 95% Confidence Intervals reported. Binary logistic regressions and multivariate linear regression models were run to determine the significance and direction of association of the paths from the predictor variable to the mediator variable, and from the mediator variable to the outcome variable.

## Results

### Bivariate analysis - engaging in weekly biking (yes/no) was significantly associated with wayfinding distance

A point-biserial correlation indicated that weekly biking yes/no were significantly associated with wayfinding distance respectively, when controlling for multiple comparisons (*p* < 0.005, bonferroni-corrected with 10 comparisons)(Table 2). All other associations were non-significant (*p* > 0.005, bonferroni-corrected with 10 comparisons)(Tables 2 and 3).

**Table 2.** Spearman’s correlations between each of the continuous variables included in the main model and weighted wayfinding distance. *Point-biserial correlations were run to measure the association between the binary variables included in the model and weighted wayfinding distance. Significant p-values are highlighted in bold.

| Variable | *r* | *p* |
| --- | --- | --- |
| Weekly biking yes/no* | 0.15 | <0.001 |
| Weekly boating yes/no* | 0.09 | 0.019 |
| Weekly public transport yes/no* | 0.08 | 0.041 |
| Age one started to drive solo | 0.08 | 0.047 |
| Average GPS reliance score | 0.07 | 0.048 |
| Hours of weekly exercise in the last 6 months | 0.05 | 0.207 |
| Hours of driving per week | 0.04 | 0.268 |
| Age one started to learn to drive | 0.03 | 0.474 |
| Hours of weekly exercise | 0.03 | 0.476 |
| Hours of walking per week | 0.01 | 0.706 |

**Table 3.**
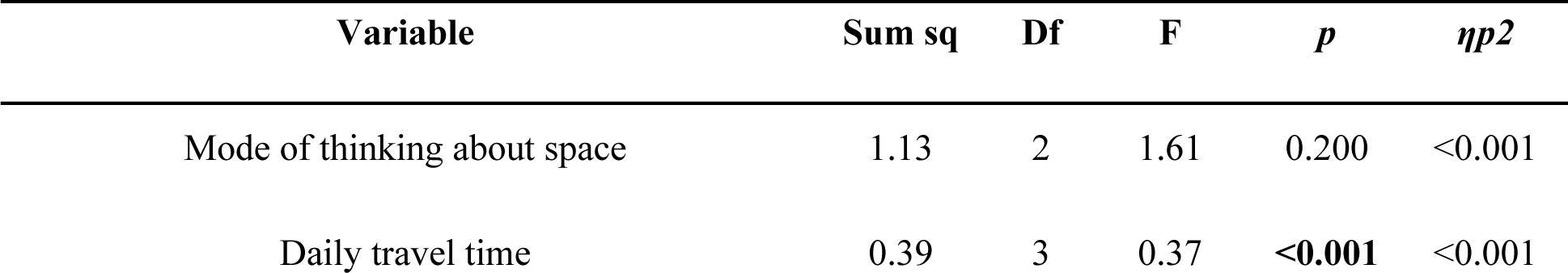
Associations between daily travel time, mode of thinking about space and weighted wayfinding distance. P-values for the significant associations are highlighted in bold.

### Multivariate analysis - the age of one started to drive solo and engaging in weekly biking (yes/no) were significantly associated with wayfinding distance, when controlling for other variables

After running these initial correlations, we then ran a linear multivariate regression model to verify these associations when controlling for the associations of other variables with wayfinding distance, with these travel-related variables as key variables of interest, and with age, gender, hours of weekly gaming on all devices and hours of weekly phone use as covariates. Interactions between gender and each of the sleep variables were included in the model (Table 2). Before running the regression, as a check for multicollinearity, we calculated the *GVIF*^[*1*/(*2*∗*df*)]^ value for each predictor variable in the model. This revealed that all the predictor variables had a *GVIF*^[*1*/(*2*∗*df*)]^ value that was less than 5, indicating that multicollinearity was not a concern (Supplementary Table S1).

**Table 4.**
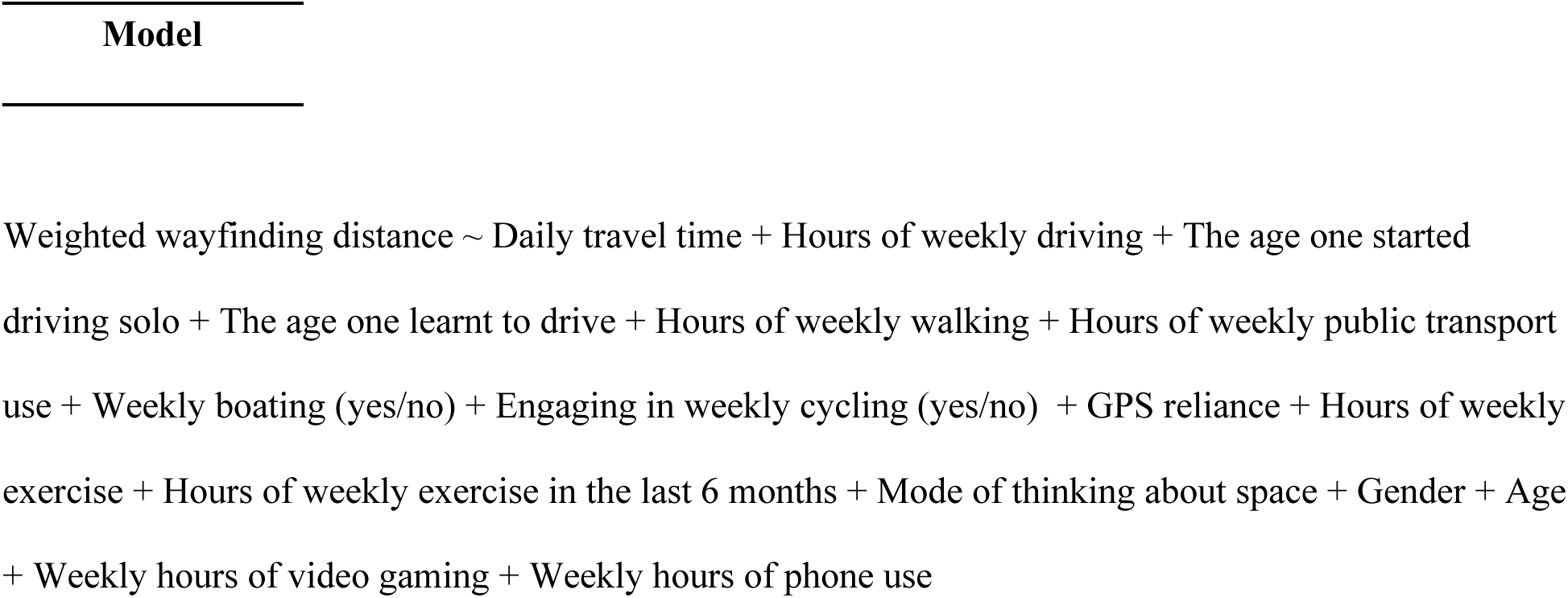
Model specification.

The outputs from the multivariate linear regression model are as follows:

### Main variables of interest

#### Age one learnt to drive, age one started driving solo and weekly hours of driving

There was a significant main effect of the age one started to drive solo on wayfinding distance (*β* = 0.11, *f2* = 0.01, *p* = 0.015, *CI* = [0.02, 0.19])(Table 5 and Figure 2). Post-hoc t-tests revealed that those who started to learn to drive solo below 18 years old were significantly better navigators than those who started to learn to drive solo aged 18 and above (*t* = -2.07, *p* = 0.039)(Table 6). Neither the age one learnt to drive (*β* = -0.06, *f2* = <0.01, *p* = 0.172, *CI* = [-0.14, 0.03]) or the weekly hours one spent driving (*β* = 0.01, *f2* = <0.01, *p* = 0.793, *CI* = [-0.04, 0.05]) were significantly associated with wayfinding distance (Table 5 and Supplementary Figure S1A-B).

**Figure 2.**
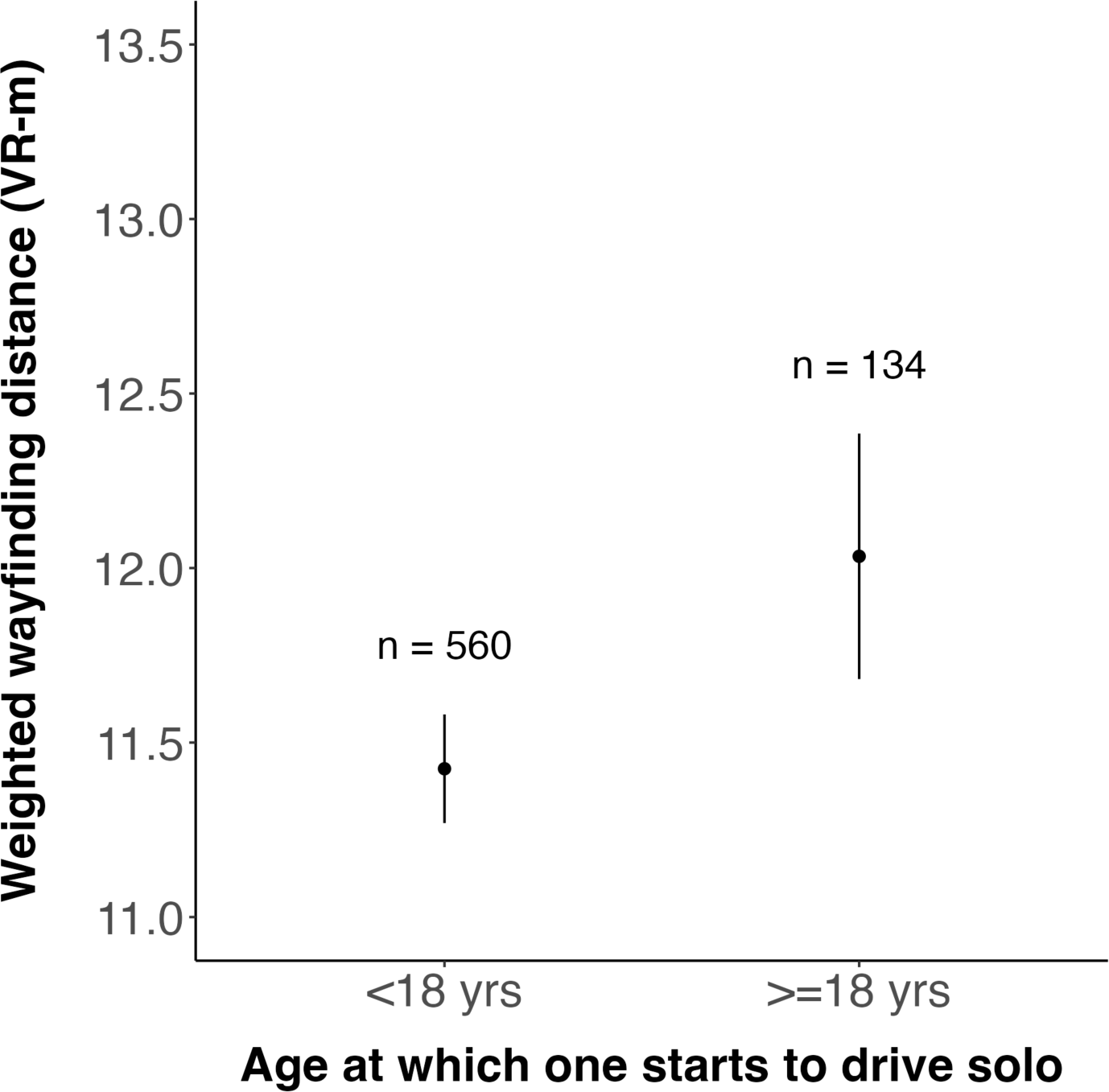
Weighted wayfinding distance in those who started to learn to drive below 18 compared with those who started to drive solo at 18 or above. VR-m = virtual reality meters. Data points represent the mean wayfinding distance across game levels across participants. Bars represent the standard error of the mean corresponding to this wayfinding distance.

**Table 5.** Model output. P-values for the significant associations are highlighted in bold.^a^ The Cohen’s f2 effect size for daily travel time was calculated for the main effect of the variable as a whole rather than for each category/level of the variable.

| Variable | $\beta$ | 95% CI | $t$ | $p$ | sig | $f^2$ |
| --- | --- | --- | --- | --- | --- | --- |
| (Intercept) | -0.12 | [-0.32, 0.08] | -1.15 | 0.250 |  |  |
| Age one learnt to drive | -0.06 | [-0.14, 0.03] | -1.37 | 0.172 |  | <0.01 |
| Age one starts to drive solo | 0.11 | [0.02, 0.19] | 2.44 | **0.015** | * | 0.01 |
| Mode of thinking about space | 0.05 | [-0.12, 0.23] | 0.60 | 0.550 |  | <0.01 |
| Weekly hours of driving | 0.01 | [-0.04, 0.05] | 0.26 | 0.793 |  | <0.01 |
| Weekly hours of exercise | -0.04 | [-0.09, 0.02] | -1.23 | 0.221 |  | <0.01 |
| Weekly hours of walking | 0.02 | [-0.03, 0.07] | 0.89 | 0.372 |  | <0.01 |
| Weekly biking - yes | 0.18 | [0.06, 0.29] | 3.02 | **0.003** | ** | 0.02 |
| Weekly boating - yes | 0.09 | [-0.16, 0.33] | 0.68 | 0.495 |  | <0.01 |
| Hours of daily travel | -0.03 | [-0.12, 0.06] | -0.59 | 0.554 |  | <0.01 |
| Weekly hours of exercise over the last 6 months | 0.08 | [-0.03, 0.19] | 1.39 | 0.165 |  | <0.01 |
| Average GPS reliance score | 0.02 | [-0.02, 0.07] | 1.11 | 0.267 |  | <0.01 |
| Age | 0.12 | [0.07, 0.17] | 5.14 | **<0.001** | *** | 0.03 |
| Gender | -0.18 | [-0.28, -0.08] | -3.50 | **0.001** | *** | 0.06 |
| Weekly hours of video gaming on all devices | -0.11 | [-0.16, -0.06] | -4.65 | **<0.001** | *** | 0.03 |
| Weekly hours of phone use | 0.02 | [-0.02, 0.07] | 0.94 | 0.346 |  | <0.01 |
| Highest level of education achieved | 0.03 | [-0.07, 0.12] | 0.54 | 0.589 |  | <0.01 |

**Table 6.** Bonferroni-corrected post-hoc t-tests comparing weighted wayfinding distance in those starting to drive solo below the age of 18 with those who started doing so at aged 18 or above. P-values for the significant associations are highlighted in bold. VR-m = virtual reality meters.

| Comparison | *t* | *p* |
| --- | --- | --- |
| <18 yrs vs >=18 yrs | -2.07 | **0.039** |

### Covariates

Please see Supplementary Figures S2A-M for visualisations of the associations between each of the covariates with wayfinding distance.

### Mediation analysis - the significant association between engaging in weekly biking (yes/no) and wayfinding distance was significantly mediated by gender

A mediation analysis was run to determine whether the significant associations between both the age one started to drive solo and whether one bikes weekly (yes/no) and wayfinding distance were mediated by age, gender, weekly hours of video game use on all devices, the type of environment one grew up in (city or not) and the type of environment one currently lives in (city or not).

The significant association between the age one started driving solo and wayfinding distance was not significantly mediated by either age, gender, gaming hours per week on all devices, whether one grew up in a city or whether one currently lives in a city, as shown by the non-significant indirect effects of these variables (*p* > 0.05)(Supplementary tables S2-S6). The significant association between whether one bikes weekly and wayfinding distance was significantly mediated by gender, as shown by a significant indirect effect of gender (*p* = 0.034)(Supplementary table S8). A binary logistic regression model showed that those who engage in weekly biking are significantly more likely to be female than those not (*β* = 0.58, *OR* = 1.78, *p* = 0.019, *CI* = [0.10, 1.06])(Supplementary table S9), where females were significantly worse navigators than males (*β* = -0.18, *f2* = 0.06, *p =* 0.001, *CI* = [-0.28, -0.08])(Table 5). The significant association between weekly biking (yes/no) and wayfinding distance was not significantly mediated by either age, gaming hours per week on all devices, whether one grew up in a city or whether one currently lives in a city, as shown by the non-significant indirect effects of these variables (*p* > 0.05)(Supplementary tables S7,S10, S11 and 12).

## Results summary

Our key finding was that the age at which one started driving solo was significantly associated with wayfinding ability, where those who start driving solo below 18 were significantly better at wayfinding than those who start driving solo at aged 18 and above, when accounting for other travel-related variables.

## Discussion

Although many studies have shown that specific facets of spatial cognition are associated with driving experience and performance (Andrews & Westerman, 2012; Di Meco et al., 2021; Kunishige et al., 2020; Moran et al., 2020; Tinella et al., 2020, 2021a, 2021b), our study was the first to investigate the relationship between self-reported driving experience and measured wayfinding ability. As hypothesised, we found a significant association between the age one started to drive solo and wayfinding distance, where those who started driving solo below 18 were significantly better at navigating than those who started driving solo aged 18 and above. This significant association was not significantly mediated by either age, gender, video game experience, the type of environment one currently lives in or the type of environment one grew up in, suggesting that these factors could not explain why those who started driving solo at a younger age were better navigators. However, we did not find a significant association between hours of weekly driving or the age one started to learn to drive with wayfinding ability.

One potential explanation for the link between learning to drive and navigation ability is that those people who learn earlier have greater spatial perspective taking, mental rotation and visuospatial processing ability which lead to being able to learn to drive faster and learning to navigate better. This is supported by evidence that better performance on different facets of spatial cognition are associated with better driving performance (Andrews & Westerman, 2012; Di Meco et al., 2021; Kunishige et al., 2020; Moran et al., 2020; Tinella et al., 2020, 2021a, 2021b) and navigation ability (Garg et al., 2023; Wolbers and Hegarty, 2010). Indeed, spatial skills can predict driving test success in clinical and non-clinical samples (Barco et al., 2014; Barrash et al., 2010; Crizzle et al., 2012). An alternative explanation is that those learning to drive solo at a younger age may use their new driving skill to spend more time exploring new places over a wider range than the others, giving them greater exposure to navigational challenges and honing navigation skill. This explanation is consistent with research suggesting exposure to different experiences at a young age can shape later life abilities (Everson-Rose, 2003; Lee & Schafer, 2020). More specifically, this explanation is consistent with our recent research using Sea Hero Quest that found a link between the street network entropy of the environment encountered growing up and later life spatial ability (Coutrot et al., 2022a). The greater sample size of those who started driving solo before 18 years old (n = 560) compared with those who started driving solo at aged 18 and above (n = 134) could be explained by the fact that those living in the US are reliant on using cars for transport in most places, making learning to drive early important for one’s independence. This would also align with the notion that exposure to different experiences in youth can shape later life abilities.

One possibility that we considered was that those who started driving alone at a younger age were participants who had grown up in rural settings, and due to the advantage conferred from rural environments on navigation skill (Coutrot et al. 2022a), had gained greater spatial skills from the environment rather than the driving. However, the non-significant mediation effect of both the environment one grew up in and currently lives in make this possibility unlikely. If one learns to drive early it would suggest they would likely have longer experience in driving, which in itself could lead to greater spatial skills. Given that we did not ask participants about the number of years of driving experience they had, we were unable to determine whether the effect of starting to drive solo at a younger age was associated with simply having a greater number of years of driving experience. However, the non-significant association between the age one learnt to drive and wayfinding ability suggests that the period of time over which one drives is unlikely to account for the significant association between the age one started to drive solo and wayfinding ability. Rather, it appears that the early exposure to driving or the aptitude to learn to drive are more important in determining navigation ability.

It is important to consider that not all studies have shown that spatial cognition is associated with driving performance. Accordingly, a recent study showed that memory and global cognitive function were significant predictors of driving performance whilst visuospatial processing was not (Ledger et al., 2019b). Other affect-related measures such as trait anger and behavioural impulsivity have also been associated with poorer self-reported and objective driving performance (Difonzo et al., 2022; Stojan et al., 2021; Yu et al., 2022). Additionally, attitudes towards driving, such as whether one enjoys driving, may have also influenced the association between the age one started to drive solo and wayfinding performance. Thus, further studies should take into consideration other cognitive and affective factors when examining the association between driving experience and navigation ability.

### Limitations and future directions

Whilst our study allowed assessment in 694 participants on an ecologically valid (Coutrot et al., 2019) and reliable cognitive test of navigation (Coughlan et al., 2020), there are a number of limitations future studies could consider and address. It would be useful to explore other variables related to driving, such as driving frequency, years of driving experience, the distance covered and the type of environment one usually drives in, using direct driving-related metrics, such as distance traveled daily and diversity of routes taken. Additionally, the participant sample we recruited using prolific may not have been representative of the general population, and so recruiting participants from a wider range of sources would help overcome this issue. Future studies would also benefit from longitudinal designs, given the reciprocal relationship between driving and cognitive function (Choi et al., 2013; Edwards et al., 2008), and use of a broader range of virtual navigation and spatial memory tests that extend to real-world environments (Brown et al., 2010; Burles & Iaria, 2020; Coutrot et al., 2019; He et al., 2021; Hegarty et al., 2006; Hill et al., 2023; Howard et al., 2014; Javadi et al., 2019; Patai et al., 2019; van der Ham et al., 2022). Critically it will be important to examine whether greater spatial cognitive skills predict learning to drive solo, or whether it is the exposure to environment made possible by driving early, or a combination, that predicts later navigation ability.

### Conclusion

Overall, our findings indicate that early life experiences, such as driving, may be important in predicting future navigation ability. Accordingly, this research will serve as a platform for future research looking into how actively engaging in such activities in early life may improve one’s ability to navigate in complex environments and provide a greater understanding as to what drives the wide variation in individual differences in navigation ability.

## Acknowledgements

We would like to thank all the participants who volunteered to take part in this research. This research is part of the Sea Hero Quest initiative funded and supported by Deutsche Telekom. Alzheimer’s Research UK (ARUK-DT2016-1) funded the analysis; Glitchers designed and produced the game; and Saatchi and Saatchi London managed its creation.

## Data availability

A dataset containing the preprocessed trajectory lengths and demographic information is available at https://osf.io/cugdw/. We also set up a portal - https://seaheroquest.alzheimersresearchuk.org/ - where researchers can invite a targeted group of participants to play SHQ and generate data about their spatial navigation capabilities. Those invited to play the game will be sent a unique participant key, generated by the SHQ system according to the criteria and requirements of a specific project decided by the experimenter. Access to the portal will be granted for non-commercial purposes. Future publications based on this dataset should add ‘Sea Hero Quest Project’ as a co-author.

## Code availability

The code used to produce this data is available at: https://osf.io/cugdw/

