## Supplementary Materials for "Individuals learning to drive solo before age 18 have superior spatial navigation ability compared with those who learn later"

| **Variable** | ***VIF*** |
| --- | --- |
| Age one learnt to drive | 4.13 |
| Age one started to drive solo | 4.22 |
| Mode of thinking about space | 1.04 |
| Weekly hours of driving | 1.26 |
| Weekly hours of exercise | 1.91 |
| Weekly hours of walking | 1.32 |
| Weekly biking (yes/no) | 1.22 |
| Weekly boating (yes/no) | 1.16 |
| Hours of daily travel | 1.18 |
| Weekly hours of exercise over the last 6 months | 1.73 |
| GPSaverage | 1.12 |
| Age | 1.23 |
| Gender | 1.41 |
| Weekly hours of video gaming on all devices | 1.25 |
| Weekly hours of phone use | 1.14 |
| Highest education level achieved | 1.12 |

Table S1. None of the predictor variables in the model had a *VIF* value exceeding 5, suggesting that multicollinearity was not a concern.

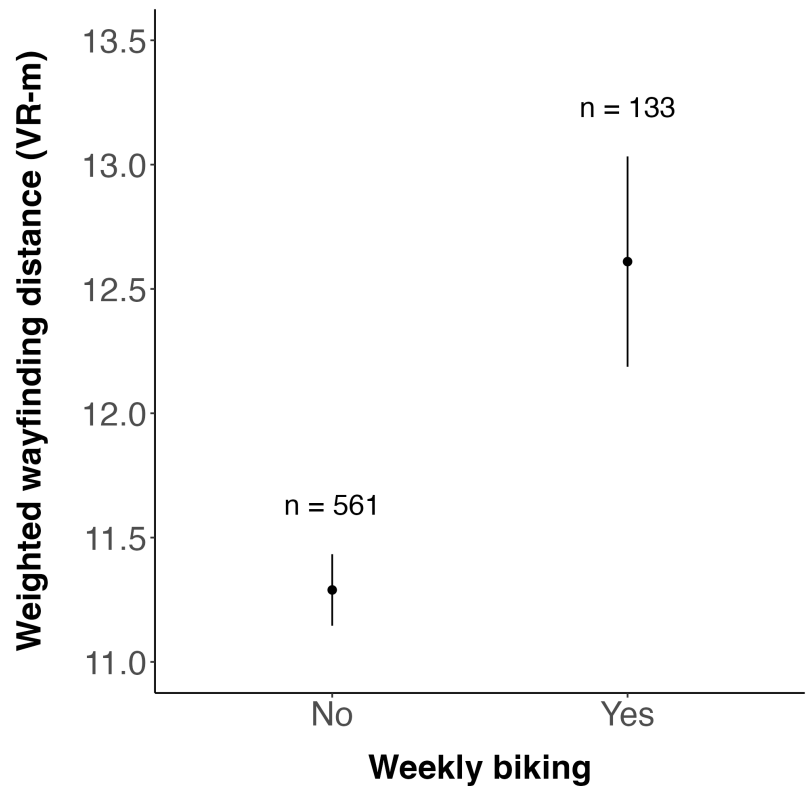

Figure S1. Post-hoc t-tests comparing wayfinding distance in those who reported biking weekly compared with those who did not. VR-m = virtual reality metres.

| Age | Estimate | 95% CI | *p* |
| --- | --- | --- | --- |
| Average casual mediation effect | <0.01 | [-0.01, 0.01] | 0.972 |
| Average direct effect | 0.05 | [0.01, 0.10] | 0.006 |
| Total effect | 0.05 | [0.01, 0.10] | 0.008 |
| Proportion mediated | <0.001 | [-0.29, 0.19] | 0.979 |

**Table S2. The association between the age one started driving solo and weighted wayfinding distance when using age as a mediator variable.**

| **Gender** | **Estimate** | **95% CI** | **p** |
| --- | --- | --- | --- |
| Average casual mediation effect | <0.01 | [-0.01, 0.01] | 0.659 |
| Average direct effect | 0.05 | [0.01, 0.10] | 0.008 |
| Total effect | 0.06 | [0.01, 0.10] | 0.007 |
| Proportion mediated | 0.04 | [-0.18, 0.24] | 0.658 |

**Table S3. The association between the age one started driving alone and weighted wayfinding distance when using gender as a mediator variable.**

| **Weekly hours of video gaming** | **Estimate** | **95% CI** | **p** |
| --- | --- | --- | --- |
| Average casual mediation effect | -0.01 | [-0.02, <0.01] | 0.242 |
| Average direct effect | 0.05 | [0.01, 0.10] | 0.008 |
| Total effect | 0.05 | [0.01, 0.09] | 0.020 |
| Proportion mediated | -0.11 | [-0.77, 0.09] | 0.262 |

**Table S4. The association between the age one started driving solo and weighted wayfinding distance when using weekly hours of video gaming as a mediator variable.**

| **Growing up in a city** | **Estimate** | **95% CI** | ***p*** |
| --- | --- | --- | --- |
| Average casual mediation effect | <0.01 | [<0.01, <0.01] | 0.697 |
| Average direct effect | 0.05 | [0.01, 0.10] | 0.008 |
| Total effect | 0.05 | [0.01, 0.10] | 0.008 |
| Proportion mediated | <0.001 | [-0.02, 0.01] | 0.694 |

**Table S5. The association between the age one started driving solo and weighted wayfinding distance when using the environment one grew up in (city or not) as a mediator variable.**

| **Currently living in a city** | **Estimate** | **95% CI** | ***p*** |
| --- | --- | --- | --- |
| Average casual mediation effect | <0.01 | [<0.01, <0.01] | 0.907 |
| Average direct effect | 0.05 | [0.01, 0.10] | 0.006 |
| Total effect | 0.05 | [0.01, 0.10] | 0.006 |
| Proportion mediated | 0.01 | [-0.06, 0.08] | 0.901 |

**Table S6. The association between the age one started driving solo and weighted wayfinding distance when using the environment one currently lives in (city or not) as a mediator variable.**

| **Age** | **Estimate** | **95% CI** | ***p*** |
| --- | --- | --- | --- |
| Average casual mediation effect | 0.01 | [-0.01, 0.05] | 0.204 |
| Average direct effect | 0.18 | [0.05, 0.33] | 0.006 |
| Total effect | 0.20 | [0.06, 0.35] | 0.004 |
| Proportion mediated | 0.07 | [-0.05, 0.29] | 0.205 |

**Table S7. The association between weekly biking (yes/no) and weighted wayfinding distance when using age as a mediator variable.**

| **Gender** | **Estimate** | **95% CI** | ***p*** |
| --- | --- | --- | --- |
| Average casual mediation effect | -0.01 | [-0.04, <0.01] | **0.040** |
| Average direct effect | 0.18 | [0.05, 0.33] | 0.008 |
| Total effect | 0.17 | [0.03, 0.31] | 0.018 |
| Proportion mediated | -0.09 | [-0.60, <0.01] | 0.058 |

**Table S8. The association between weekly biking (yes/no) and weighted wayfinding distance when using gender as a mediator variable.**

| **Variable** | **$\beta$** | **95% CI** | ***t*** | ***p*** | **sig** | **OR** |
| --- | --- | --- | --- | --- | --- | --- |
| (Intercept) | -1.20 | [-2.12, -0.32] | -2.62 |  |  |  |
| Age one learnt to drive | 0.27 | [-0.08, 0.65] | 1.48 | 0.139 |  | 1.32 |
| Age one starts to drive solo | -0.13 | [-0.50, 0.24] | -0.67 | 0.505 |  | 0.88 |
| Mode of thinking about space | 0.77 | [-0.01, 1.58] | 1.91 | 0.056 | . | 2.16 |
| Weekly hours of driving | 0.04 | [-0.16, 0.25] | 0.42 | 0.674 |  | 1.04 |
| Weekly hours of exercise | 0.27 | [0.02, 0.52] | 2.14 | **0.033** | * | 1.31 |
| Weekly hours of walking | -0.15 | [-0.38, 0.06] | -1.37 | 0.170 |  | 0.86 |
| Weekly biking - yes | 0.58 | [0.10, 1.06] | 2.35 | **0.019** | * | 1.78 |
| Weekly boating - yes | 0.28 | [-0.78, 1.36] | 0.51 | 0.611 |  | 1.32 |
| Hours of daily travel | 0.30 | [-0.10, 0.70] | 1.46 | 0.146 |  | 1.35 |
| Weekly hours of exercise over the last 6 months | 0.29 | [-0.18, 0.76] | 1.21 | 0.225 |  | 1.34 |
| Average GPS reliance score | -0.28 | [-0.47, -0.09] | -2.91 | **0.004** | ** | 0.76 |
| Gender | 0.37 | [0.18, 0.56] | 3.80 | **<0.001** | *** | 1.45 |
| Weekly hours of video gaming on all devices | 0.98 | [0.79, 1.19] | 9.61 | **<0.001** | *** | 2.67 |
| Weekly hours of phone use | -0.50 | [-0.71, -0.29] | -4.64 | **<0.001** | *** | 0.61 |
| Highest education level achieved | -0.55 | [-0.97, -0.13] | -2.58 | **0.010** | ** | 0.58 |

**Table S9. Binary logistic regression model predicting gender using weekly biking (yes/no) and other travel and navigation-related covariates. P-values for the significant associations are highlighted in bold. OR = Odds Ratio.**

| **Weekly hours of video gaming** | **Estimate** | **95% CI** | **p** |
| --- | --- | --- | --- |
| Average casual mediation effect | 0.01 | [-0.01, 0.03] | 0.232 |
| Average direct effect | 0.18 | [0.04, 0.33] | 0.008 |
| Total effect | 0.19 | [0.05, 0.34] | 0.007 |
| Proportion mediated | 0.06 | [-0.06, 0.22] | 0.235 |

**Table S10. The association between weekly biking (yes/no) and weighted wayfinding distance when using weekly hours of video gaming as a mediator variable.**

| **Growing up in a city** | **Estimate** | **95% CI** | ***p*** |
| --- | --- | --- | --- |
| Average casual mediation effect | <0.01 | [-0.01, 0.01] | 0.958 |
| Average direct effect | 0.18 | [0.05, 0.34] | 0.008 |
| Total effect | 0.18 | [0.05, 0.34] | 0.008 |
| Proportion mediated | <0.01 | [-0.05, 0.04] | 0.960 |

**Table S11. The association between weekly biking (yes/no) and weighted wayfinding distance when using the environment one grew up in (city or not) as a mediator variable.**

| **Currently living in a city** | **Estimate** | **95% CI** | ***p*** |
| --- | --- | --- | --- |
| Average casual mediation effect | <0.01 | [-0.02, 0.01] | 0.444 |
| Average direct effect | 0.19 | [0.05, 0.33] | 0.008 |
| Total effect | 0.18 | [0.05, 0.33] | 0.010 |
| Proportion mediated | -0.01 | [-0.15, 0.05] | 0.452 |

**Table S12. The association between weekly biking (yes/no) and weighted wayfinding distance when using the environment one currently lives in (city or not) as a mediator variable.**

| Comparison | *t* | *p* |
| --- | --- | --- |
| >=18 and <21 yrs vs >=21 and <24 yrs | -0.14 | 0.890 |
| >=18 and <21 yrs vs >=24 and <30 yrs | -0.94 | 0.349 |
| >=18 and <21 yrs vs >= 30 yrs | -3.78 | **<0.001** |
| >=21 and <24 yrs vs >=24 and <30 yrs | -0.82 | 0.410 |
| >=21 and <24 yrs vs >= 30 yrs | -3.77 | **<0.001** |
| >=24 and <30 yrs vs >= 30 yrs | -3.14 | **0.002** |

**Table S13. Bonferroni-corrected post-hoc t-tests for the main effects of age on weighted wayfinding distance. P-values for the significant associations are highlighted in bold when applying a bonferroni-corrected alpha threshold of 0.017.**

| Comparison | *t* | *p* |
| --- | --- | --- |
| Men vs women | -4.66 | **<0.001** |

**Table S14. Post-hoc t-tests for the main effects of gender on weighted wayfinding distance. P-values for the significant associations are highlighted in bold when applying a an alpha threshold of 0.05.**

| Comparison | *t* | *p* |
| --- | --- | --- |
| >=1 and <5 hpw vs >=5 and <10 hpw | 3.04 | **0.002** |
| >=1 and <5 hpw vs >= 10 hpw | 5.38 | **<0.001** |

|  |  |  |
| --- | --- | --- |
| $\geq 5$ and $< 10$ hpw vs $\geq 10$ hpw | 2.17 | 0.031 |

**Table S15. Bonferroni-corrected post-hoc t-tests for the main effects of weekly video gaming on weighted wayfinding distance. P-values for the significant associations are highlighted in bold when applying a bonferroni-corrected alpha threshold of 0.008.**

| Comparison | *t* | *p* |
| --- | --- | --- |
| $\geq 1$ and $< 10$ hpw vs $\geq 10$ and $< 20$ hpw | -0.92 | 0.358 |
| $\geq 1$ and $< 10$ hpw vs $\geq 20$ and $< 30$ hpw | -0.57 | 0.567 |
| $\geq 1$ and $< 10$ how vs $\geq 30$ and $< 40$ hpw | -1.28 | 0.200 |
| $\geq 1$ and $< 10$ hpw vs $\geq 40$ hpw | -0.96 | 0.337 |
| $\geq 10$ and $< 20$ hpw vs $\geq 20$ and $< 30$ hpw | 0.46 | 0.646 |
| $\geq 10$ and $< 20$ hpw vs $\geq 30$ and $< 40$ hpw | -0.46 | 0.643 |
| $\geq 10$ and $< 20$ hpw vs $\geq 40$ hpw | 0.07 | 0.948 |
| $\geq 20$ and $< 30$ hpw vs $\geq 30$ and $< 40$ hpw | -0.93 | 0.355 |
| $\geq 20$ and $< 30$ hpw vs $\geq 40$ hpw | -0.44 | 0.660 |
| $\geq 30$ and $< 40$ hpw vs $\geq 40$ hpw | 0.57 | 0.567 |

**Table S16. Bonferroni-corrected post-hoc t-tests for the main effects of hours of weekly phone use on navigation performance when applying a bonferroni-corrected alpha threshold of 0.005.**

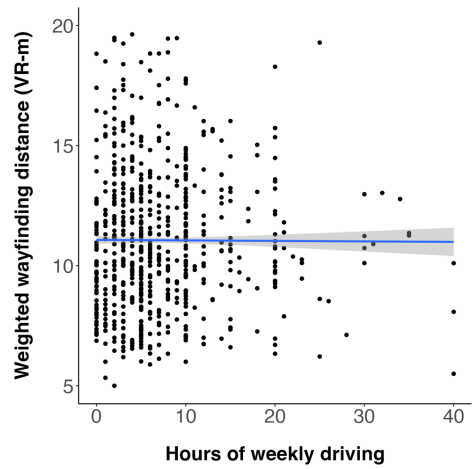

A)

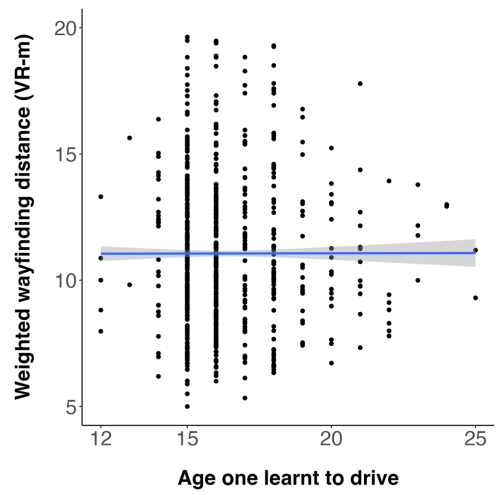

B)

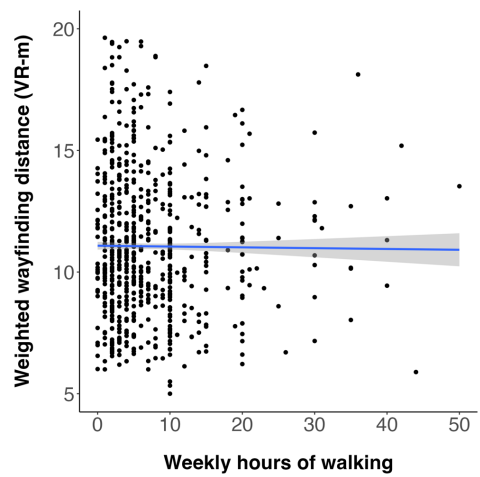

C)

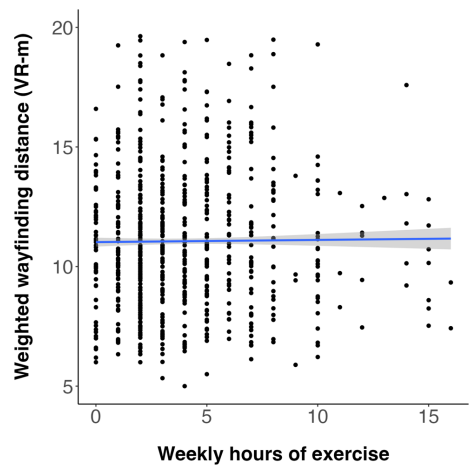

D)

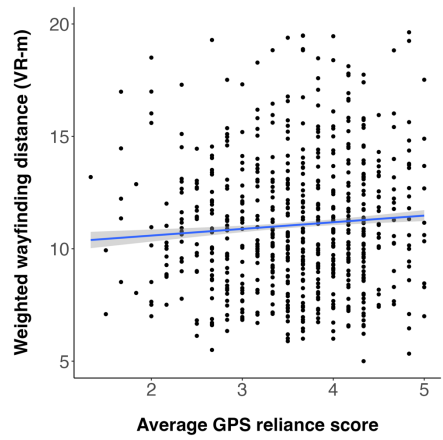

E)

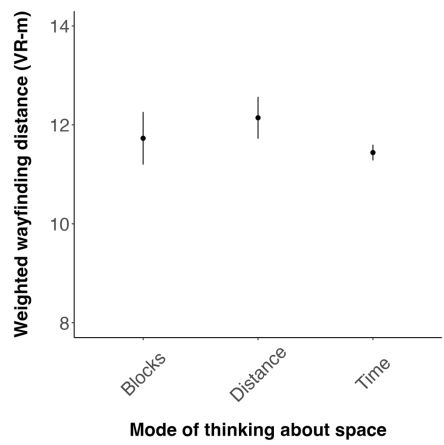

F)

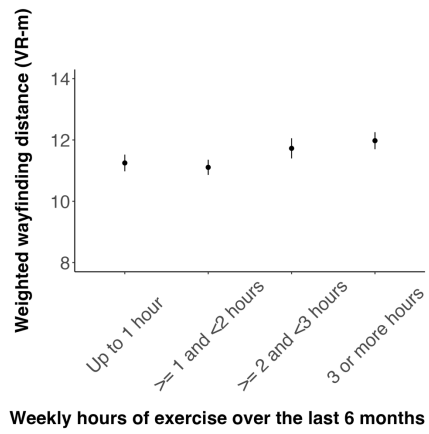

G)

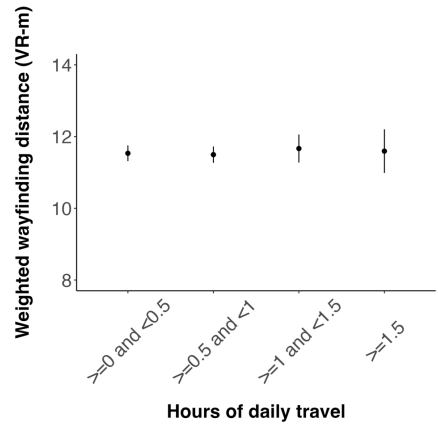

H)

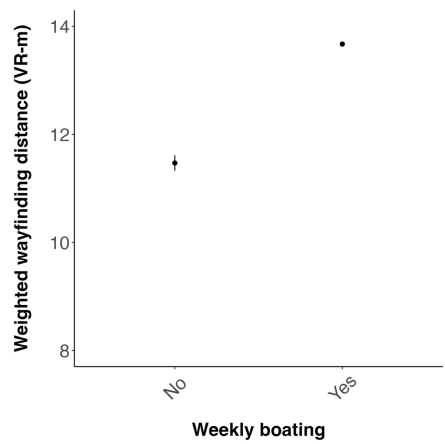

I)

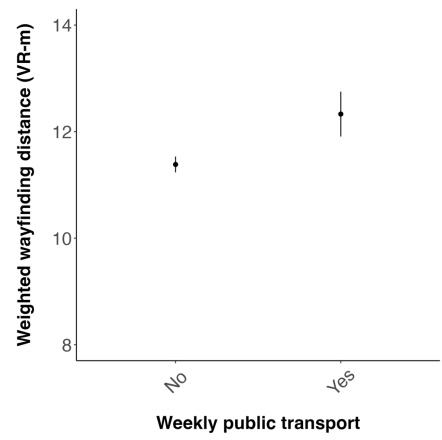

J)

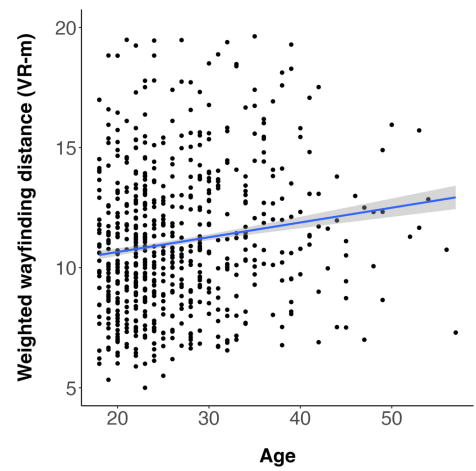

L)

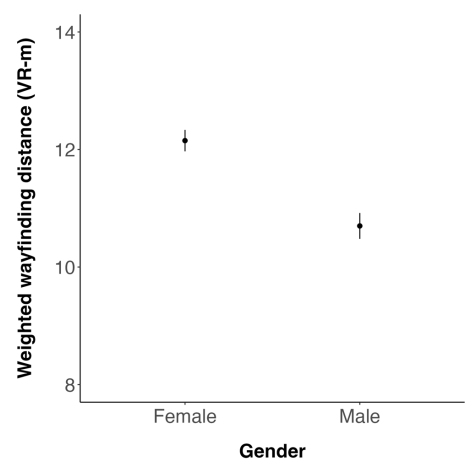

M)

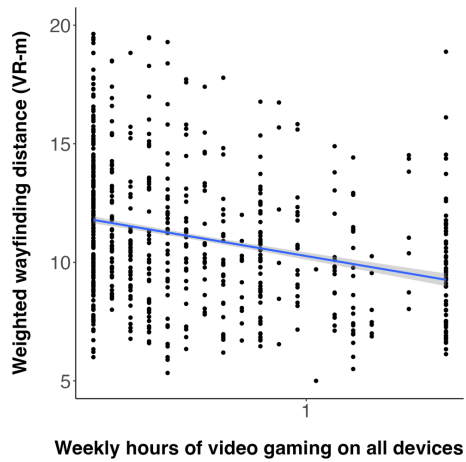

N)

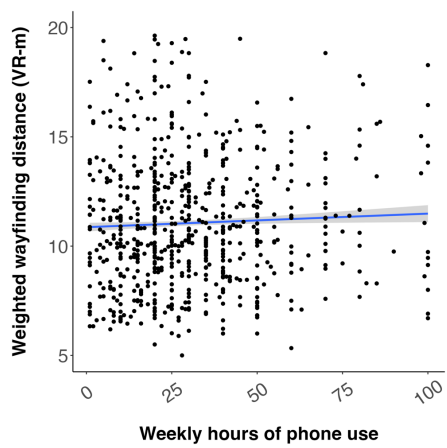

O)

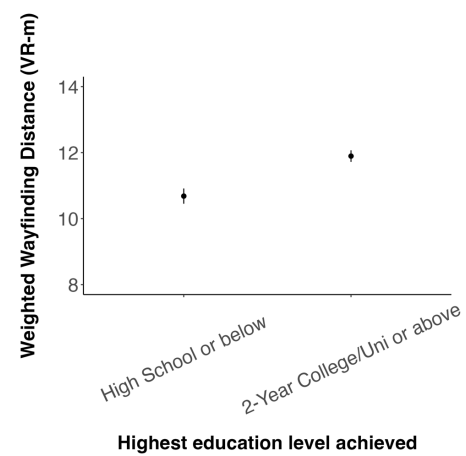

P)

**Figure S2. (A-P) Associations between each of the travel-related and demographic variables and weighted wayfinding distance.**

**(A-E,L,N-O) Blue line indicates the mean wayfinding distance across game levels across participants. Grey shading surrounding the blue line indicates the standard error of the mean corresponding to this wayfinding distance. Data points indicate the mean wayfinding distance across game levels for an individual participant.**

**(F-K,M,P) Data points represent the mean wayfinding distance across game levels across participants. Bars represent the standard error of the mean corresponding to this wayfinding distance.**

#### Questionnaires

##### **Demographics:**

1. How old are you?
2. What gender are you? (male, female, other)
3. What is the highest level of education you have achieved? (no formal education, some formal education, high school, 2-year college/university, 4-year college/university, masters, PhD)

##### **Gaming experience:**

1. How often do you play video games per week? (0 = minimum, 20+ hours = maximum)
2. How often do you use a smartphone or a tablet per week? (0 = minimum, 40+ hours = maximum)

### **GPS Reliance Scale (Dahmani and Bohbot, 2020):**

**\*(question 5 was omitted from the main analysis)**

For all questions below, 1 = never, 2 = sometimes, 3 = often, 4 = very often, 5 = always.

1. How often would you use GPS to navigate in general?
2. You are meeting friends at a new restaurant, and you are travelling there for the first time. How often do you use a GPS in such a situation?
3. You usually leave from home to go to a doctor's appointment. This time, however, your appointment is scheduled right after work. Therefore, you have to travel a new route to get to a destination you have visited before. How often do you use a GPS in such a situation?
4. You usually leave from home to visit your family. You are taking the same route as you always do. How often do you use a GPS in such a situation?
5. You usually travel a specific route to go to your friend's house. This time, you think you may get there faster by taking a different route. How often do you take new routes to travel to places you have visited before?
6. How often do you use a GPS to travel to a destination outside of the area where you live?
7. When finding your way around a city outside of your hometown, how often do you use a GPS?

### **Travel-related questions:**

1. How many hours per week do you walk? (0 = min, 100+ = max)
2. How many hours per week do you ride a bike? (0 = min, 100+ = max)
3. How many hours per week do you drive a car/taxi? (0 = min, 100+ = max)
4. How many hours per week do you use public transport? (0 = min, 100+ = max)

5. How many hours per week do you navigate on a boat? (0 = min, 100+ = max)
6. What is your daily travel time? (0-0.5 hours = min, 30 minute intervals, 5.5-6 hours = max)
7. How physically active were you in the last 6 months? (up to one hour per week, 1-2 hours per week, 2-3 hours per week, 3 or more hours per week, compete professionally)
8. How many hours of exercise do you perform in an average week? (0 = min, 20+ = max)
9. What age did you learn to drive? (0 = min, 65 = max, if you did not learn to drive, please leave the slider at 0)
10. What age did you start driving alone? (0 = min, 65 = max, if you have never driven, please leave the slider at 0)
